# Interneuronal gap junctions increase synchrony and robustness of hippocampal ripple oscillations

**DOI:** 10.1101/311662

**Authors:** André Holzbecher, Richard Kempter

## Abstract

Sharp wave-ripple (SWRs) are important for memory consolidation. Their signature in the hippocampal extracellular field potential (EFP) can be decomposed into a ≈ 100 ms long sharp wave superimposed by ≈ 200 Hz ripple oscillations. How ripple oscillations are generated is currently not well understood. A promising model for the genesis of ripple oscillations is based on recurrent interneuronal networks (INT-INT). According to this hypothesis, the INT-INT network in CA1 receives a burst of excitation from CA3 that generates the sharp wave, and recurrent inhibition leads to an ultrafast synchronization of the CA1 network causing the ripple oscillations; fast-spiking parvalbumin-positive basket cells (PV+BCs) may constitute the ripple-generating interneuronal network. PV+BCs are also coupled by gap junctions (GJs) but the function of GJs for ripple oscillations has not been quantified. Using simulations of CA1 hippocampal networks of PV+BCs, we show that GJs promote synchrony and increase the neuronal firing rate of the interneuronal ensemble, while the ripple frequency is only affected mildly. The promoting effect of GJs on ripple oscillations depends on fast GJ transmission (≲ 0.5 ms), which requires proximal gap junction coupling (≲ 100 μm from soma).

## 2 Introduction

Sharp wave-ripple (SWRs) are transients of the extracellular field potential (EFP) in the hippocampus that occur in rest and slow-wave sleep (Buzsáki et al., 1992). SWRs consist of ≈ 200 Hz ripple oscillations that are enveloped by the ≈ 100 ms long sharp wave (O’Keefe, 1976; O'Keefe and Nadel, 1978). Different computational tasks are associated with SWRs, e.g., decision making via preplaying future trajectories (Diba and Buzsáki, 2007) or memory consolidation (Girardeau et al., 2009) putatively via replay of past events (Wilson and McNaughton, 1994), which underlines their role for the two-stage model of memory consolidation (Buzsáki, 1989).

The underlying mechanism that is responsible for generating ripples is debated (Traub et al., 2000; Memmesheimer, 2010; Ylinen et al., 1995). Three distinct origins of ripples were proposed, which could also coexist: (1) the axonal plexus of pyramidal neurons, which is putatively densely coupled by GJs (Draguhn et al., 1998; Traub et al., 1999), (2) supralinear synaptic integration in feedforward excitatory networks (Memmesheimer, 2010; Jahnke et al., 2015) and (3) fast recurrent interneuronal networks (INT-INT, Ylinen et al., 1995). In silico, all of these models can create ripple-like oscillations.

Recent experiments support the INT-INT hypothesis. In CA3, Schlingloff et al. (2014) showed in vitro that optogenetic activation of PV+ interneurons leads to ripple-like oscillations, even with excitatory chemical transmission blocked. This matches well the results from an in vivo study by Stark et al. (2014), where blocking of the interneurons abolished ripple oscillations in CA1. Following this hypothesis, evidence accumulates that from the zoo of interneurons (Chamberland and Topolnik, 2012) fast-spiking parvalbumin-expressing basket cells (PV+BCs) are a key component of the INT-INT ripple-generating network (Klausberger et al., 2003; Schlingloff et al., 2014). PV+BCs are highly active during SWRs and fire phase-locked to ripple oscillations (Klausberger et al., 2003). Furthermore, INT-INT networks can explain intraripple frequency accommodation (Donoso et al., 2018), an experimental hallmark of SWRs (Ponomarenko et al., 2004).

Additional to the strong and fast GABAergic synapses, PV+BCs are coupled by gap junctions (GJs; Fukuda and Kosaka, 2000b; Tamás et al., 2000; Galarreta and Hestrin, 2001a,b; Bartos et al., 2002). Two major categories of experiments exists to determine the function of GJs in SWRs (review Posłuszny, 2014): pharmacological GJ blockers (Ylinen et al., 1995) and genetic knockouts of GJ proteins (Hormuzdi et al., 2001; Güldenagel et al., 2001). However, the experimental results prove ambiguous and hence inconclusive, i.e., many of the experiments that use GJ blockers find a strong suppression of SWRs (Ylinen et al., 1995; Draguhn et al., 1998; Hormuzdi et al., 2001; Maier et al., 2003; Pais et al., 2003; Traub et al., 2003; Buhl et al., 2003; Behrens et al., 2011), while the studies relying on GJ knockout mice find a rather mild effect (Hormuzdi et al., 2001; Pais et al., 2003; Buhl et al., 2003; Behrens et al., 2011).

In addition to experiments, there are also substantial contributions from theoretical neuroscience to understand the function of GJs in oscillations. These results were derived by both analytical (Lewis and Rinzel, 2003; Pfeuty et al., 2005; Ostojic et al., 2009; Tchumatchenko and Clopath, 2014) and numerical means (Traub et al., 2001; Maex and De Schutter, 2003; Bartos et al., 2007; Guo et al., 2012; Fink et al., 2015).

On the analytical side, studies from Lewis and Rinzel (2003) and Pfeuty et al. (2005) of two-neuron systems have shown that the effect of GJs on synchrony depends on the proportions of electrical and chemical coupling. Using mean-field analysis, Ostojic et al. (2009) showed that oscillations can arise in a neuronal network that is exclusively coupled by GJs but the network frequency equals the mean firing rate of the neurons.

While these studies provided a sound theoretical basis, they were not specifically tailored to reproduce biologically realistic networks. Numerical simulations give more insights on this issue, e.g., GJs were found to increase synchrony in gamma oscillations (30–70 Hz, Traub et al. 2001; Bartos et al. 2007; Guo et al. 2012) and ripple oscillations (Maex and De Schutter, 2003). However, Maex and De Schutter restricted their analysis to one set of GJ parameters, which does not satisfy the variability found in the GJ connectivity data.

Even though there is evidence that PV+BCs are coupled by GJs and that PV+BCs are the generator of ripples, the implications of interneuronal GJ-coupling for ripple oscillations have not been investigated in depth. Thus, here we study how interneuronal GJs of different coupling probabilities, conductances, and delays impact the primary properties of hippocampal ripple oscillations, which advances our understanding about this phenomenon.

## 3 Methods

Since we want to investigate the influence of interneuronal GJs on hippocampal ripple oscillations, we first present the network model that we use for our analysis. Subsequently, we introduce the measures used to characterize ripple oscillations. Finally, we present the multicompartment models of hippocampal PV+BCs that are used for measuring the amplitudes and the delays of the GJ coupling potentials between two neurons. A summary of all standard parameters is given in Table 1, and an overview of all the varied parameters for each figure is given in Table 2.

**Table 1:** Standard parameters for all simulation except stated otherwise. If the “Methods” are referenced, a further explanation for the value of the parameter is given therein.

| Name | Symbol | Value | Reference |
| --- | --- | --- | --- |
| **Gap junctions** |  |  |  |
| Active-spike component | $\beta$ | 0.25 mV | Galarreta and Hestrin, 2001a |
| Passive conductance | $\gamma$ | 1 nS | Galarreta and Hestrin, 2001a |
| Multicompartment GJ conductance | $g^{\text{GJ}}$ | 1 nS | Galarreta and Hestrin, 2001a |
| **Inhibitory synapses** |  |  |  |
| Latency | $\tau_l$ | 1.0 ms | Bartos et al., 2002 |
| Rise time | $\tau_r$ | 0.45 ms | Bartos et al., 2002 |
| Decay time | $\tau_d$ | 1.2 ms | Bartos et al., 2002 |
| Peak conductance | $g^{\text{peak}}$ | 5 nS | Bartos et al., 2002 |
| Reversal potential | $v^{\text{inh}}$ | -75.0 mV | Buhl et al., 1995 |
| **Excitatory synapses** |  |  |  |
| Latency |  | 1 ms | Taxidis et al., 2012 |
| Rise time |  | 0.5 ms | Taxidis et al., 2012 |
| Decay time |  | 2 ms | Taxidis et al., 2012 |
| Peak conductance |  | 1 nS | Taxidis et al., 2012 |
| Reversal potential | $v^{\text{exc}}$ | 0 mV | Taxidis et al., 2012 |
| **Membrane properties of PV+BCs** |  |  |  |
| AP threshold |  | -52 mV | Wang and Buzsáki, 1996 |
| Reset potential |  | -67 mV | Wang and Buzsáki, 1996 |
| Refractory period |  | 1.0 ms | Wang and Buzsáki, 1996 |
| Leak reversal potential | $v^{\text{leak}}$ | -65.0 mV | Buhl et al., 1996 |
| Leak conductance | $g^{\text{leak}}$ | 10 nS | Buhl et al., 1996 |
| Capacitance | $C$ | 100 pF | Buhl et al., 1996 |
| Membrane time constant |  | 10 ms | Buhl et al., 1996 |
| **Network properties** |  |  |  |
| Network size |  | 200 | Donoso et al., 2018 |
| Overall GJ connectivity | $p_{\text{GJ}}$ | 0.06 | Methods |
| GJ connectivity to nearest neighbor |  | 0.3 | Methods |
| GJ nearest neighbors |  | 40 | Methods |
| Inhibitory connectivity |  | 0.2 | Donoso et al., 2018 |
| Averaged drive per neuron |  | 4000 spikes/s | Donoso et al., 2018 |

**Table 2:**
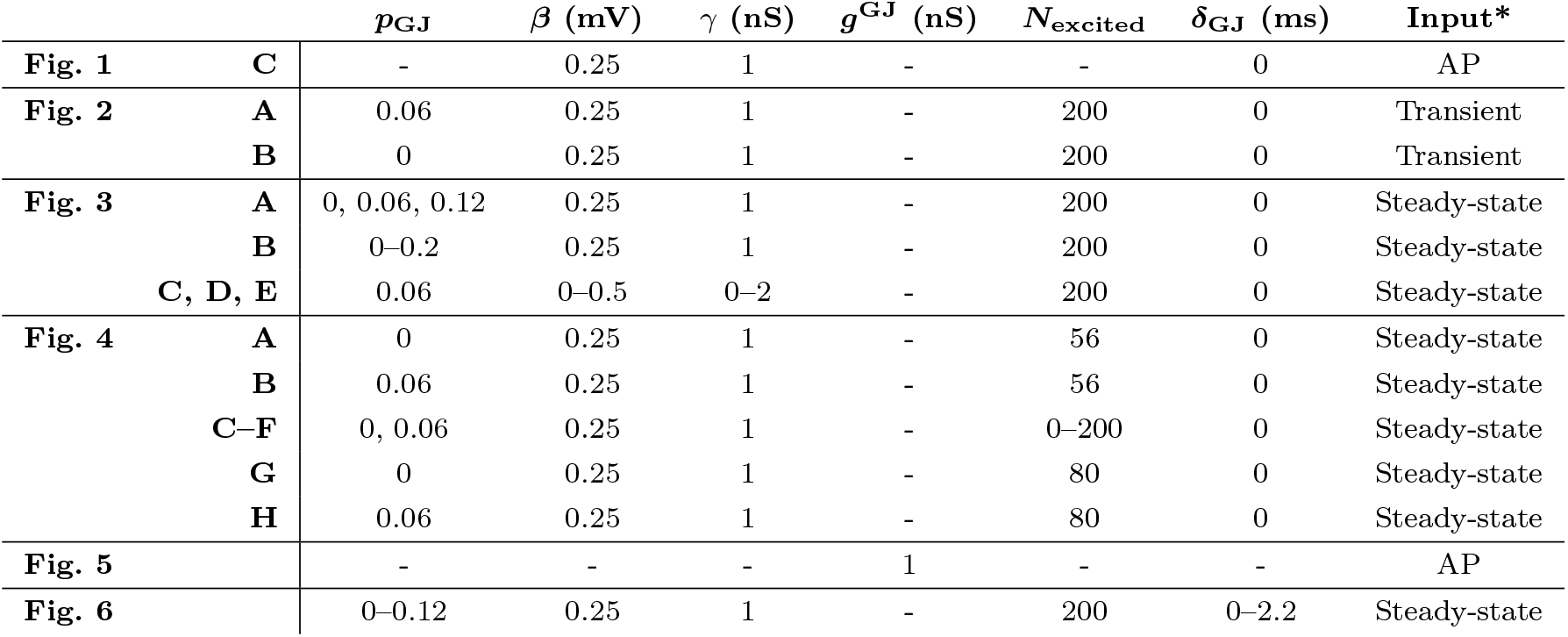
Parameters varied across figures. Note that *g*^GJ^ is only used in the multicompartment model. (*) Input to the neuron or the interneuronal network. Here, “AP” stands for a sufficient input to stimulate an action potential; “Transient” for transient sharp wave-like excitation at a peak rate of ≈ 2000 spikes/s and a background Poisson rate of 750 spikes/s; “Steady-state” for Poisson steady-state input at 4000 spikes/s.

| | | $p_{\text{GJ}}$ | $\beta$ (mV) | $\gamma$ (nS) | $g^{\text{GJ}}$ (nS) | $N_{\text{excited}}$ | $\delta_{\text{GJ}}$ (ms) | Input* |
| --- | --- | --- | --- | --- | --- | --- | --- | --- |
| **Fig. 1** | **C** | - | 0.25 | 1 | - | - | 0 | AP |
| **Fig. 2** | **A** | 0.06 | 0.25 | 1 | - | 200 | 0 | Transient |
|  | **B** | 0 | 0.25 | 1 | - | 200 | 0 | Transient |
| **Fig. 3** | **A** | 0, 0.06, 0.12 | 0.25 | 1 | - | 200 | 0 | Steady-state |
|  | **B** | 0–0.2 | 0.25 | 1 | - | 200 | 0 | Steady-state |
|  | **C, D, E** | 0.06 | 0–0.5 | 0–2 | - | 200 | 0 | Steady-state |
| **Fig. 4** | **A** | 0 | 0.25 | 1 | - | 56 | 0 | Steady-state |
|  | **B** | 0.06 | 0.25 | 1 | - | 56 | 0 | Steady-state |
|  | **C–F** | 0, 0.06 | 0.25 | 1 | - | 0–200 | 0 | Steady-state |
|  | **G** | 0 | 0.25 | 1 | - | 80 | 0 | Steady-state |
|  | **H** | 0.06 | 0.25 | 1 | - | 80 | 0 | Steady-state |
| **Fig. 5** |  | - | - | - | 1 | - | - | AP |
| **Fig. 6** |  | 0–0.12 | 0.25 | 1 | - | 200 | 0–2.2 | Steady-state |

### CA1 network model

Hippocampal PV+BCs are highly active during SWRs and fire phase locked to ripple oscillations. Thus, we consider a minimal model for ripple generation in the CA1 hippocampal area that only consists of PV+BCs, and neglects all other neuron types as motivated by Donoso et al. (2018).

In this network model, the neurons are approximated by point-like leaky integrate-and-fire (LIF) neurons to set the focus on the network dynamics. In total, we simulate 200 neurons, which resembles the average number of PV+BCs in an in vitro slice preparation of the hippocampal area CA1 (Nimmrich et al., 2005; Donoso et al., 2018). The interneurons are coupled by gap junctions and inhibitory synapses, and they receive Poisson distributed excitation (Fig. 1A).

**Figure 1:**
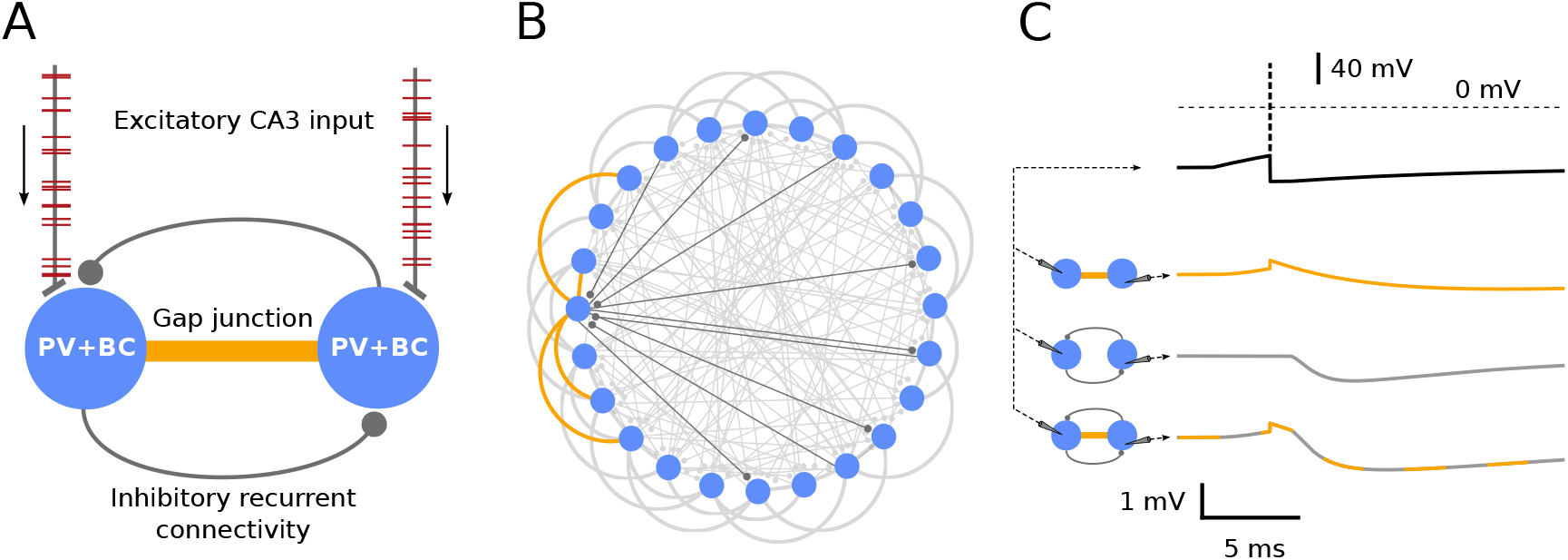
Network architecture. **A**, Network schematic of the CA1 PV+BC network that receives Poisson distributed excitation representing inputs from the CA3 region (arrows). The CA1 neurons are coupled by gap junctions (orange) and inhibitory synapses (gray lines with circles at their ends). **B**, More detailed scheme of connectivity. To resemble an in-vitro slice preparation, the network is chosen to consist of 200 neurons (here only 24 shown). Gap junctions are introduced with a connectivity of 0.3 between the 20% nearest neighbors, i.e., 0.06 overall connectivity (curves, highlighted in orange for one neuron on the left). Interneurons are randomly connected with a probability of 0.2 by inhibitory synapses (gray lines; highlighted in dark gray for one neuron on the left). **C**, Membrane potentials of a postsynaptic neuron show the response to a presynaptic action potential (top, black) when neurons are connected by only a gap junction (orange), only an inhibitory synapse (gray) or both (orange/gray). Standard parameters of all simulations are displayed in Tables 1 and 2.

### Neuron and synapse model

The dynamics of the membrane potential *v_i_* of neuron *i* is modeled by

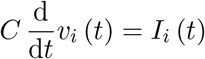

for *v_i_* that is below the firing threshold of −52.0 mV (Wang and Buzsáki, 1996). When the membrane voltage reaches this threshold the neuron fires an action potential (AP). Subsequently, the neuron is reset to −67.0 mV and remains inactive for a refractory period of 1.0 ms (Wang and Buzsáki, 1996). The capacitance is set to *C* = 100 pF for all neurons (Buhl et al., 1996). Further, the current *I_i_* (*t*) that is received by the *i*th neuron is given by

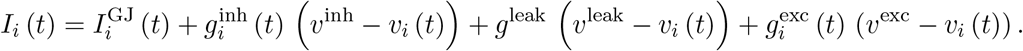

Here *g*^leak^ = 10 nS and *v*^leak^ = −65.0 mV (Buhl et al., 1996) are the leak conductance and the leak reversal potential, respectively. The analogous functions and parameters for inhibition and excitation are 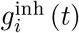 and *v*^inh^ = −75.0 mV (Buhl et al., 1995), and 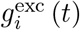 and *v*^exc^ = 0 mV (Taxidis et al., 2012), respectively. The membrane time constant set by this leak conductance and the membrane capacitance is 10 ms (Buhl et al., 1996).

The gap junction current 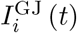 is realized following an approach from Lewis and Rinzel (2003) and Ostojic et al. (2009). In this model, the current transferred through the gap junction to neuron *i* from all GJ coupled neurons *j* is given as (Fig. 1C)

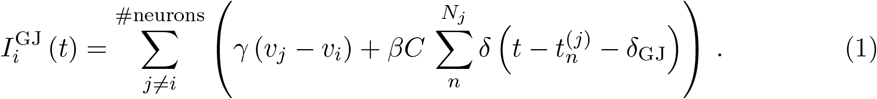

Here, the first term in the sum accounts for the passive subthreshold coupling. The bidirectional current is assumed to be ohmic, and hence given by the multiplication of the GJ conductance *γ* and the membrane potential difference of cell *j* and cell *i*. Since LIF neurons do not model the spiking dynamics of the membrane potential, the “postsynaptic” potential caused by a “presynaptic” action potential is introduced manually by the second term. Naturally, gap junctions are bidirectional, and “postsynaptic” refers to the neuron, which receives a GJ potential and elicits no spike. Analogously, “presynaptic” is used hereafter to describe the neuron that elicited an action potential. Here, *β* is the amount of voltage added to the membrane potential of the postsynaptic neuron *i* at each presynaptic spiking event *n* (*n* = 1, …, *N_j_*) at times 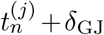 of presynaptic neuron *j*. For this to hold true, the neuronal capacitance *C* is introduced in the expression, so that the product of *C* and *β* has the dimension of a charge. The parameter *δ*_GJ_ delays the response of the postsynaptic neuron to a presynaptic action potential and accounts for dendritic latencies. Since the active-spike component *β* mediates fast potentials, and hence more sensitive to a delay, the delay is only included in the active-spike component *β*. Biologically realistic values for *β* and *γ* can be extracted from electrophysiological studies (Galarreta and Hestrin, 1999; Tamás et al., 2000; Galarreta and Hestrin, 2001a), and here we use *β* = 0.25 mV and *γ* = 1.0 nS in most simulations. When we explore the effects of *β* and *γ* on the network dynamics, *β* is varied in the range [0, …, 0.5] mV and *γ* in the range [0, …, 2.0] nS.

The inhibitory GABAergic conductances are modeled by a biexponential function (Fig. 1C)

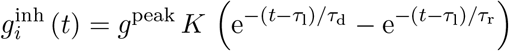

for a spike event at *t* = 0 and *t* > *τ*_1_, otherwise 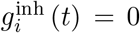. Here, *τ*_1_ = 1.0 ms sets the latency till the onset of the response, *τ*_r_ = 0.45 ms is the rise time constant, and *τ*_d_ = 1.2 ms is the decay time constant of the conductance. The peak conductance is given by *g*^peak^ = 5 nS, and *K* is a normalization chosen such that the maximal value of the 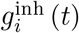 is 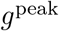 (Bartos et al., 2002).

Moreover, each neuron receives an excitatory Poisson-like input, which is mediated by the time dependent conductance 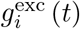. Here, we use the same model as for the inhibitory synapses but with *g*^peak^ = 1 nS, *τ*_1_ = 1.0 ms, *τ*_r_ = 0.5 ms, and *τ*_d_ = 2.0 ms (Taxidis et al., 2012).

### Network connectivity

There are in total three types of different synaptic connections in the network model: feedforward excitation, recurrent inhibition, and bidirectional gap junctional coupling. Here, we briefly motivate the connectivities used in our model.

Most of the studies of gap junctional coupling between PV+BC investigated neocortical areas (Tamás et al., 2000; Galarreta and Hestrin, 1999, 2001a; Amitai et al., 2002). They found that interneurons that are further than 200 μm apart are only rarely coupled by GJs since for dendritic GJ coupling the dendritic fields of the neurons have to overlap. However, within a radius of 200 μm the values found for the connection probability of PV+BCs are high: 59 % (Amitai et al., 2002), 61% (Gibson et al., 1999) and 66 % (Galarreta and Hestrin, 1999). Data for hippocampal networks is less abundant. The GJ connection probability found in dentate gyrus varies from 29 % (Bartos et al., 2001) to up to 92 % (Hormuzdi et al., 2001). Data for the hippocampal area CA3 is provided by Hormuzdi et al. (2001) and Bartos et al. (2002), who found 5 of 8 (63%) and 3 of 6 (50%) fast spiking, parvalbumin-positive neuron pairs to be GJ coupled, respectively. Bartos et al. (2002) found in CA1 that 2 of 9 basket cell pairs were electrically coupled. Further evidence for GJ coupling in the area CA1 comes from ultrastructural studies (Katsumaru et al., 1988; Fukuda and Kosaka, 2000b), which show the existence of GJ coupling, however electrophysiological studies that quantify GJ coupling in CA1 in more detail are, to our knowledge, not available (Bartos et al., 2002).

In conclusion, we set the standard connection probability for neurons within a distance of 200 μm to 30 %, and to 0 % otherwise. Note that in our network model a spatial structure is only introduced by GJs, which couple neurons to their nearer neighbors (Fig. 1B). When we explore the influence of the GJ connection probability on the network dynamics, we vary the connection probability in the range from 0 to 100 %.

The number of nearest neighbors in a vicinity of 200 μm is around 40 interneurons, since the total extent of the ventral hippocampal CA1 slice is around 1.1 × 0.4 × 0.1 mm^3^ (Dougherty et al., 2012). For this approximation, we assume that the 200 neurons of the model are distributed homogeneously in space, but we neglect the 0.1 mm width of the pyramidal cell layer. Moreover, we take into account the effective size of the dendritic field in the 0.4 mm direction of the slice that is reduced by cutting the slices, i.e., most of the dendritic trees will not lie completely within the slice.

When we combine the number of neurons that are within a sphere of 200 μm with the assumed connectivity of 30 %, we find the overall GJ connectivity in the network to be *p*_GJ_ = 0.06 (Fig. 1B), i.e., one neuron is coupled to 12 other neurons via GJs on average.

Furthermore, the interneurons are coupled by random recurrent inhibition with 20 % connectivity according to estimates by Donoso et al. (2018) for ventral hippocampal slices. Excitation, of which 10 % is shared, is fed into the interneurons in form of Poisson distributed spike trains at 4000 spikes/s per interneuron in the simulations of the steady-state dynamics. In the case of the transient dynamics, each neuron receives 35 APs, whose times are drawn from a Gaussian distribution (width of 7 ms) to model a sharp wave-like excitation (Fig. 2). At the peak of the transient excitation the rate is ≈ 2000 spikes/s.

**Figure 2:**
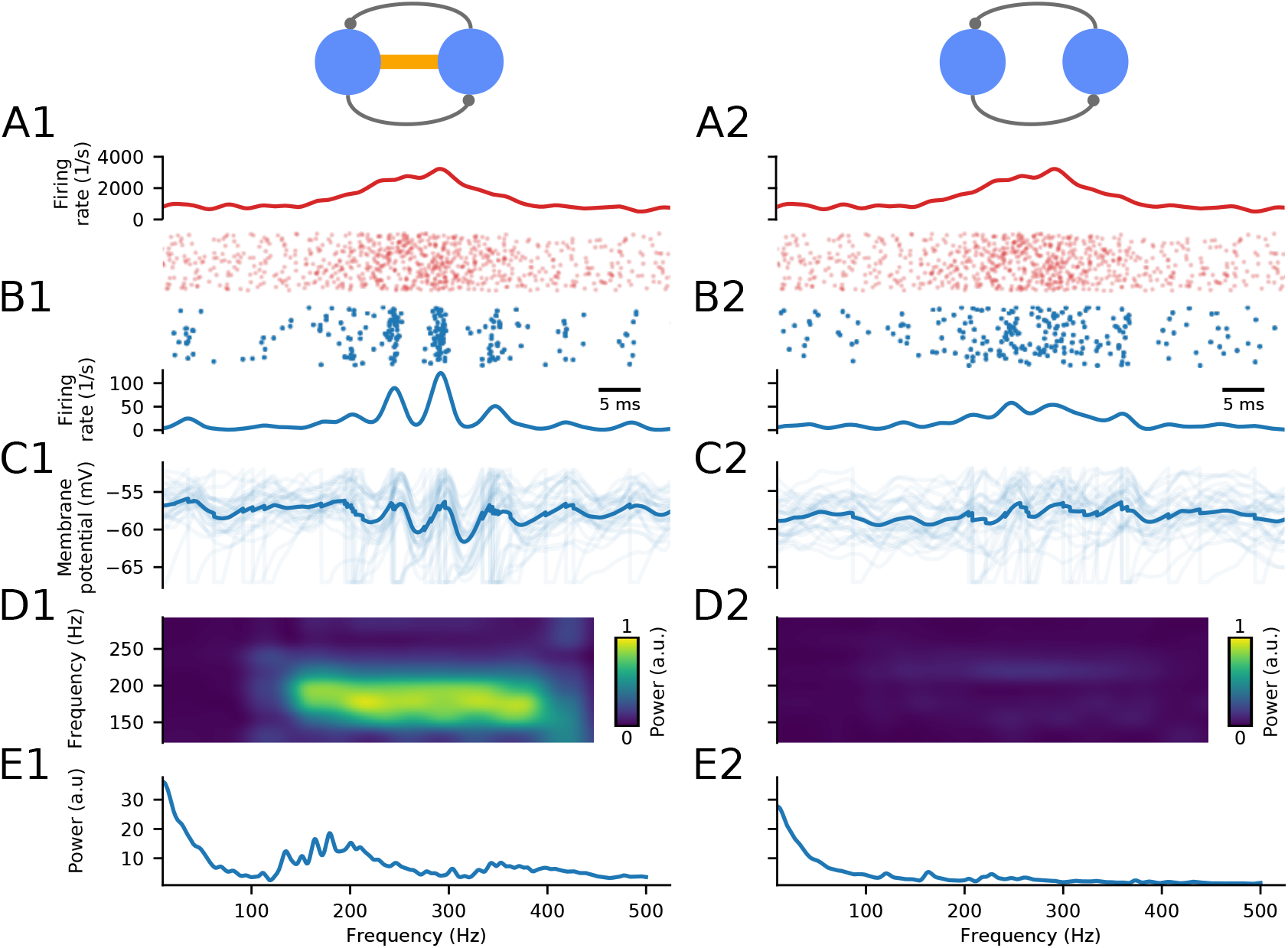
Gap junction (GJ) coupling promotes ripple oscillations during transient excitation. Ripple oscillations are much stronger in an interneuronal network (*N* = 200) with GJs (left column; *p*_GJ_ = 0.06) than in a network without GJs (right column; *p*_GJ_ = 0). **A**, Excitatory input to the interneuronal network. APs of the total excitatory population are shown in a rastergram, and average excitatory firing rates received by one interneuron (smoothed by a Gaussian filter with 1 ms width) are depicted in the plot above. Note that A–D share a common time scale. **B**, Same quantities as A but for the response of the interneuron network. **C**, Membrane potentials of interneurons. Dark blue lines represent the population average, lighter blue lines correspond to individual neurons. **D**, Spectrogram of the population activity showing elevated activity for the GJ network at ≈ 200 Hz. **E**, Power spectrum. For an overview of the parameters see Tables 1 and 2.

### Simulation routine

The simulation time is 1 s for the steady state simulations, and 0.3 s for the transient SWR oscillation. At the start of each simulation, all neurons are initialized at a random membrane potential between reset and threshold voltage. For each network simulation the connectivities of inhibition and gap junctions are set randomly.

The network simulations are carried out using the spiking network simulator “Brian” (Goodman and Brette, 2008) and provenance is ensured by using pypet (Meyer and Obermayer, 2016).

### How to characterize the network oscillation?

Measures that we use to quantify the simulated neuronal activity are: (1) the firing rate, (2) the network frequency, (3) the oscillation strength, and (4) the synchrony index.

The firing rate denotes how many times a neuron spikes per second, and it is computed over the whole time of the simulation, and averaged over the whole neuronal ensemble.

For the calculation of the network frequency, we combine the binary spike trains of the single neurons to one network spike train. Consequently, the network frequency is given by the frequency at the maximal power spectral density, i.e., the Fourier transformation of the autocorrelation function. To ignore the low-frequency components of the spectrum, we consider only frequencies > 30 Hz.

The oscillation strength is computed from the peak of the power spectral density as the product of the amplitude and the FWHM of that peak. This gives an estimate of the area under the peak that corresponds to how much power is distributed in the frequency modes around the network frequency.

The synchrony index we use is based on the pairwise event synchronization measure proposed by Quian Quiroga et al. (2002) and refined by Kreuz et al. (2007, 2015). For a spike *i* from spike train *t*^(1)^ the coincidence with a (non-empty) second spike train *t*^(2)^ is calculated by taking the minimum of the distances to each spike *j* in *t*^(2)^ and comparing it with a coincidence window *τ_c_*

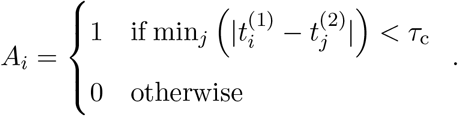

Here min_*j*_ means that the minimum is calculated over all the spikes of *t*^(2)^. In contrast to (Kreuz et al., 2007), we use a fixed coincidence window *τ*_c_ = 0.5 ms. The coincidence indicator *A_i_* is calculated for every spike *i* of every neuron. Then, all the *A_i_*’s are summed and divided by the total number of spikes *N* in all spike trains to obtain the synchrony index, and a such defined synchrony index would be between 0 and 1. To correct for spike events that are coincident by chance, which is determined by the product of 2*τ*_c_ and the average firing rate *f*, is subtracted. This yields for the synchrony index *S*

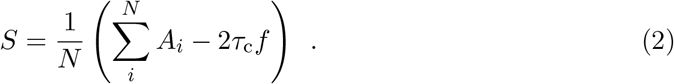

The synchrony index is a measure that is used to compare the synchrony in different configurations of the network. Accordingly, the window size of 0.5 ms is chosen such that the synchrony index leads to well distinguishable values for oscillations at ripple frequency.

For the simulation of the transient SWRs, we calculate the time-resolved spectrograms of the network activity additionally to the periodogram. This analysis is carried out in Python using the inbuilt spectrogram function from the module SciPy (Jones et al., 2001–).

### Multicompartment models for determining the GJ transmission delay

Since there is no experimental data for the delays of GJ coupling potentials in hippocampal area CA1, we use multicompartment models of hippocampal PV+BCs to calculate these. For an estimate of the properties of the GJ coupling potentials, it is important that the shape of the action potential is as close as possible to the one in real PV+BCs. Using realistic spiking dynamics as a criterion, we selected two models with a simplified basket cell morphology (Lee et al., 2014; Saudargiene et al., 2015) from ModelDB (Mc-Dougal et al., 2017; Fig. 5A). Note that the model from Saudargiene et al. (2015) has a morphology similar to the model from Lee et al. (2014) but is scaled up by roughly a factor of two (Fig. 5A). The FWHMs of the PV+BC action potential that are measured in experiments are around 0.3 ms (Buhl et al., 1996; Kohus et al., 2016) while the FWHMs for the two models are ≈ 0.6 ms (Lee et al., 2014; Saudargiene et al., 2015). The impact of this mismatch between the widths of the action potentials is discussed further in the Results.

We used the multicompartment models for calculating three different characteristics of the GJ coupling potentials. First, the delay from the maximum of the presynaptic action potential to the maximum of the postsynaptic GJ potential, i.e., the peak delay (Fig.5B left). Second, the delay from the maximal rise of the action potential to the maximal rise of the GJ coupling potential, i.e., the maximal-rise delay (Fig.5C left). And third, the amplitude of the postsynaptic GJ coupling potential (Fig.5D left). All these quantities are measured in dependence upon the soma–GJ distance. The GJ is modeled as a resistor coupling the two neurons with a conductivity of 1 nS (Galarreta and Hestrin, 2001b). For simplicity, all GJs are positioned symmetrically in the two coupled neurons, i.e., in the same dendrite and with the same distance to the soma in the respective neuron (Fig. 5A).

The three quantities, the peak delay, the maximal-rise delay, and the GJ amplitude, are estimated from the simulations, i.e., determined from the simulated voltage traces. The two coupled neurons are initialized at their equilibrium potentials, and then a current is injected in the presynaptic neuron until it elicits an action potential. More specifically, we inject a current of 0.95 nA for a duration of 4 ms into the model from Lee et al. (2014), and a current of 0.5 nA for 4 ms into the model from Saudargiene et al. (2015). These currents are chosen such that the action potential has a smooth onset. The membrane potentials of both somata are recorded while the action potential propagates from the pre-to the postsynaptic neuron.

Since the action potentials of the neuron models are too wide, we repeat the same analysis but with the AP replaced by a short, bipolar current pulse that resembles a (very) fast AP. This current pulse is generated by injecting a positive current, 0.1 ms long with an amplitude of 30 nA, and a negative current, 0.6 ms long and amplitude of −5.25 nA, into the soma of the model from Saudargiene et al. (2015), and currents of the same durations but amplitudes of 45 nA and −7.875 nA into the Lee et al. (2014) model. The bipolar current pulses are chosen such that the peak of the membrane potentials has approximately the same amplitude as an AP and the same value for both current pulses, a small afterhyperpolarization, and a minimal width. Note that for this analysis, all active conductances were switched off.

## 4 Results

The central question of our work is how gap junctions (GJs) between interneurons influence hippocampal ripple oscillations. We follow the hypothesis that hippocampal ripple oscillations are generated by recurrently connected interneurons (INT-INT), in particular, by parvalbumin-positive basket cells (PV+BCs) (Ylinen et al., 1995; Klausberger et al., 2003). Morphological and electrophysiological evidence suggests that PV+BCs are coupled by GJs (Katsumaru et al., 1988; Galarreta and Hestrin, 2001a) but their importance for ripple oscillations has not been analyzed in detail. As depicted in Fig. 1, we use a network model of 200 leaky integrate-and-fire neurons that are tuned to reproduce PV+BCs characteristics to simulate ripple oscillations as observed in acute hippocampal slice preparations (Nimmrich et al., 2005; Donoso et al., 2018).

### 4.1 Interneuronal gap junctions increase synchrony of ripple oscillations during sharp wave-like activation

To demonstrate that the PV+BC network model generates SWR-like events, the PV+BC network is first stimulated with a transient sharp wave-like input (Fig. 2). This input is modeled by a Gaussian burst of excitation (half width 7 ms, peak rate ≈ 2000 spikes/s) that resembles the excitatory inputs from area CA3 (see Methods). To this transient burst of activity a homogeneous Poisson input at 750 spikes/s is added (Fig. 2A).

Because we are interested in the difference in the network activity caused by GJs, we contrast the network dynamics of the GJ coupled network (hereafter named GJ network; Fig. 2A1–E1; GJ connection probability *p*_GJ_ = 0.06), with the network that is lacking GJ coupling (hereafter named GJ-free network; Fig. 2A2–E2; *p*_GJ_ = 0). In this example, we find that the GJ network generates ripple-like oscillations, while the GJ free network does not.

For the GJ network (Fig. 2A1–E1), we find that the neuronal population synchronizes rapidly, and collective oscillations emerge within the interneuronal population (Fig. 2B1) as a response to the Gaussian burst of excitatory input (Fig. 2A1). These oscillations are also visible in the membrane potentials of the neuronal ensemble (Fig. 2C1). The simultaneous spiking of a large fraction of the interneurons causes coincident inhibition, which leads to a strong transient hyperpolarization of the membrane potentials. The spectrogram (Fig. 2D1) shows that during the oscillation the maximal spectral power is at ≈ 200Hz, i.e., in the biologically realistic range for ripple oscillations (Maier et al., 2003). The spectrum of the full simulated network activity is shown in Fig. 2E1 to allow for a more quantitative estimate of the spectral composition of the network activity.

In contrast, in the GJ-free network, the sharp wave-like excitation does not evoke prominent ripple oscillations (Fig. 2B2&C2). The spectral analysis reveals that there are some elevated frequency modes around 180–200 Hz (Fig. 2D2&E2) but with an amplitude that is much lower than in the GJ network.

In summary, the GJ network in Fig. 2 generates prominent oscillations in the ripple frequency range whereas oscillations are weak in the GJ-free network. In these example networks, GJ parameters were set to reasonable values (Table 2). However, the electrophysiological parameters of GJs coupling of PV+BCs in hippocampal area CA1 are largely unknown (see Methods), and are object to natural variability. Thus, a more thorough analysis of a wider range of GJ parameters is necessary to account for this undetermined variability, and to test the putative role of GJs in ripple oscillations.

### 4.2 Interneuronal gap junctions synchronize steady-state ripple oscillations

To get a more quantitative estimate for the effect of GJs on the ripple oscillations generated by the CA1 network model, we explore in Fig. 3 the influence of the GJ connection probability *p*_GJ_ (for details on *p*_GJ_ see Fig. 1B). GJs can be further characterized by their active-spike component *β* and their passive conductance *γ* (Fig. 1C, and Eq. (1)). The active-spike component *β* models the amount of voltage that is added to the post-synaptic membrane potential at each presynaptic spike, and the passive conductance *γ* describes the ohmic subthreshold coupling.

**Figure 3:**
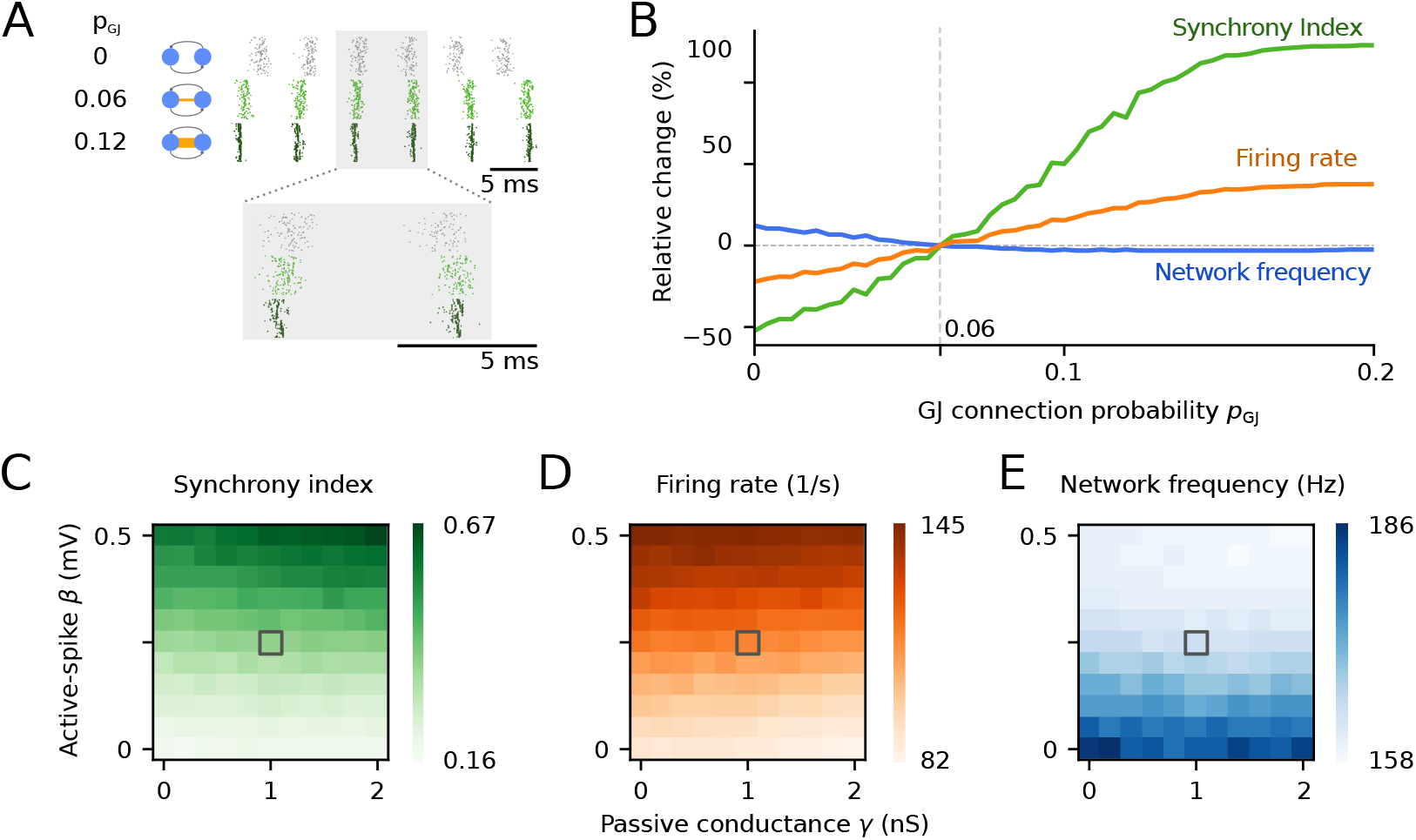
Gap junctions (GJs) increase synchrony and firing rates during ripple oscillations in interneuron networks. **A**, Rastergrams show the activities of networks without GJs (*p*_GJ_ = 0, gray, top), with standard GJ connectivity (*p*_GJ_ = 0.06; light green; middle), and with strong GJ connectivity (*p*_GJ_ = 0.12; dark green; bottom). Neurons receive Poisson input with a constant mean (4000 spikes/s). GJ parameters are *β* = 0.25 mV and *γ* = 1.0 nS. **B**, The relative change of the synchrony index, the average firing rate, and the network frequency is depicted for different values of the GJ connection probability *p*_GJ_. Quantities are normalized by their respective values at *p*_GJ_ = 0.06. GJ parameters as in A. **C–E**, GJs parameters *β* and *γ* contribute differentially to the network dynamics for fixed *p*_GJ_ = 0.06. **C**, Synchrony index as function of *β* and *γ*. **D**, Firing rate. **E**, Network frequency. Gray squares denote the GJ standard parameters as used in A and B. For an overview of the parameters see Tables 1 and 2.

To explore how *p*_GJ_, *β*, and *γ* affect ripple oscillations, we reduce the complexity of the transient network activity and analyze the network in its steady state. For example, if the network receives Poisson input at 4000 spikes/s, it oscillates at ripple frequencies (Fig. 3A) that are similar to the transient state (Fig. 2). Due to the longer temporal extent of the neuronal activity, the analysis of the network activity is more precise compared to the transient state. Simulations by Donoso et al. (2018) suggest that the results that we obtain from the steady-state ripple oscillations are transferable to transient SWR oscillations.

Examples of the steady state spiking activities of the interneuron network are shown in Fig. 3A at GJ connection probabilities *p*_GJ_ = 0, *p*_GJ_ = 0.06 (standard parameter), and *p*_GJ_ = 0.12. The networks are oscillating at 183, 163, and 159 Hz with mean interneuron firing rates of 90, 115, and 142 spikes/s for *p*_GJ_ = 0, *p*_GJ_ = 0.06, and *p*_GJ_ = 0.12, respectively. This indicates that GJs decrease the network frequency and increase the average firing rate.

In Fig. 3B, the GJ connectivity *p*_GJ_ is varied systematically, and we compute its impact on three basic network properties: the network frequency, the average firing rate, and the synchrony index. The synchrony index is a pairwise measure between 0 and 1 that counts the coincident events (coincidence window = 0.5 ms) in the neuronal spike trains; see also Methods. Simulation results are shown as the relative change of network properties with respect to their reference value at standard parameters (*p*_GJ_ = 0.06, *β* = 0.25 mV, *γ* = 1.0 nS; see Methods). The synchrony index shows the strongest dependence on the GJ connectivity. We find that an increased number of GJs in the network leads to more coincident spikes; despite a temporally compressed period of spiking activity in each oscillation cycle, also the firing rate is increased, i.e., more neurons are recruited in each cycle. Interestingly, the network frequency shows only a rather small decrease for increasing GJ connectivity (*p*_GJ_ ∈ [0, …, 0.2]).

In Fig. 3C–E, we vary the two GJ parameters *β* and *γ* independently [even though the two GJ parameters are correlated (Lewis and Rinzel, 2003; Ostojic et al., 2009)], to disentangle their effects on the synchrony index (Fig. 3C), the firing rate (Fig. 3D), and the network frequency (Fig. 3E). For all three quantities, we find that the active-spike parameter *β* has a strong influence whereas the passive parameter *γ* has only mild effects.

The synchrony index increases with increasing *β* (Fig. 3C). The same trend holds true for *γ*, however, the increase is much less pronounced. The synchrony index reaches its maximum for the maximal values of GJ parameters at *β* = 0.5 mV and *γ* = 2 nS, which is at the corner of the investigated parameter range. The average firing rate of the neuronal population also increases with increasing *β* but slightly decreases with increasing *γ* (Fig. 3D). Finally, the network frequency decreases with increasing values of *β*, and it is mildly reduced by increasing values of *γ* (Fig. 3E).

In summary, we find that introducing GJ coupling into our model network increases the synchrony and the neuronal firing rates, whereas the network frequency decreases mildly. Our simulations show that from the two GJ parameters, which describe the GJ currents (Eq. (1); Lewis and Rinzel (2003); Ostojic et al. (2009)), the active-spike component *β* is mainly responsible for the effects of the GJs on the network dynamics. The active-spike component increases the synchrony and the firing rates because it effectively acts as a precisely timed excitation that is fed into the neuronal population at the oscillation phase in which the network is on average close to spiking threshold. This increase of the interneuron firing rates leads in turn to a decrease of the network frequency because more inhibitory currents are fed back into the network. Consequently, the hyperpolarization of the membrane potentials following an oscillatory phase of spiking is stronger, hence the population needs a longer time to recover to spiking threshold, i.e., the network frequency is decreased.

For large values of *β*, the network frequency is similar to the firing rate of the neurons, i.e., every neuron is firing in almost every oscillation cycle. Rephrased in the terms introduced by Brunel (2000), this means that the increase of *β* corresponds to a transition from a synchronous irregular regime to a synchronous regular regime. Our results are in agreement with results from Ostojic et al. (2009) who found that networks that are exclusively coupled by GJs can only globally oscillate at the firing rate of the single neurons.

### 4.3 Interneuronal gap junctions reduce the number of neurons required for ripple oscillations

So far, the number of interneurons that received excitation was kept constant. However, it is not known how many interneurons are recruited during ripple oscillations and how many are required. Stark et al. (2014) optogenetically excited pyramidal cells in hippocampal networks in vivo and estimated that there are ≈ 80 pyramidal cells and ≈ 20 interneurons within the volume illuminated by the light source. Upon optic stimulation of the pyramidal cells, Stark et al. (2014) could observe fast oscillations despite the small number of pyramidal cells. However, if they optogenetically excited the ≈ 20 interneurons they could not observe oscillatory activity. In a similar experiment in vitro, Schlingloff et al. (2014) optogenetically excited ≈ 150 interneurons and showed that this number is sufficient to generate ripple-like oscillations.

Motivated by these experiments, we analyze the effect of a partial activation of the network. Therefore, the same Poisson input that was used for computing the steady-state activity in Fig. 3 is now fed into a fraction of the population of the in total 200 interneurons. The response of the interneuron network is then characterized by four properties: oscillation strength, network frequency, synchrony index, and firing rate. Moreover, we compare the dynamics of the GJ and the GJ-free network (Fig. 4).

**Figure 4:**
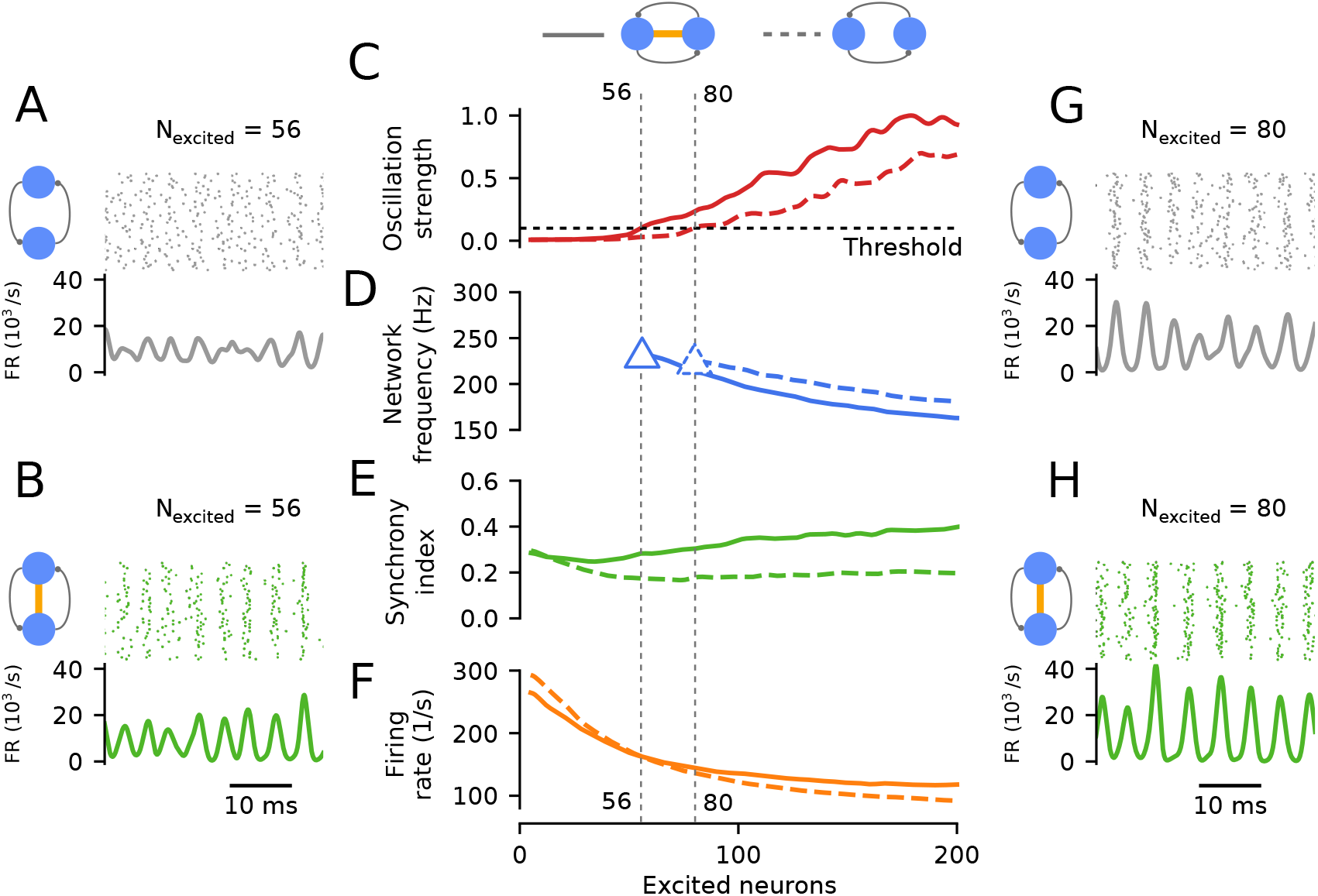
Interneuronal gap junctions (GJs) decrease the minimal number of neurons required for ripple oscillations. **A**, Example of spiking activity and network firing rate of a GJ-free network when only 56 neurons out of 200 receive excitation: oscillations are weak and unsteady. **B**, Same as A, but including GJs: oscillations are stronger and reliable. **C**, Oscillation strength as a function of the number of excited neurons of the GJ network (solid line), and of the GJ-free network (dashed line, see Methods for details). GJs decrease the number of neurons that is required to reach a certain threshold (here 0.1) of the oscillation strength. Note that **C–F** share their x-axes. **D**, Network frequency, displayed for suprathreshold oscillation strengths (solid triangle for GJ network, dashed triangle for GJ-free network). **E**, Synchrony index. **F**, Firing rate. **G,H**, Identical to A,B but for 80 excited interneurons, which is sufficient for the GJ-free network to reach the oscillation strength threshold. Plots in C–F are smoothed by a Gaussian function with a width of 5 “excited neurons”. For an overview of the parameters see Tables 1 and 2.

Four examples of the network activities with 56 and 80 excited neurons for the GJ vs. the GJ-free network are shown in Fig. 4A, B, G, H. These examples already illustrate the general trend: First, the GJ networks show stronger oscillations; second, the more neurons receive excitatory input the stronger the oscillation. Additionally, we find that only excited neurons are spiking during these simulations.

These observations are first quantified by computing the oscillation strength, which is a measure for the size of a peak in the power spectral density (see Methods for details). We introduce this measure here to be able to quantify how strong a putative network oscillation is. A reliable estimate of the network frequency is possible only if the oscillation strength is above a certain threshold (here arbitrarily chosen as 0.1; results do not critically depend on this value).

Figure 4C shows the oscillation strength as a function of the number of driven interneurons. If the oscillation strength is above the depicted threshold (0.1), oscillations are generated reliably and we consider the activity to be oscillatory, and otherwise not. The threshold is reached for the GJ network at 56 active neurons, and for the GJ-free network at 80 active neurons. In Fig. 4D, the network frequency is displayed for suprathreshold oscillation strength. In both networks, for values larger than the threshold, the network frequency is decreasing from ≈ 220 Hz to ≈ 170 Hz with increasing number of active neurons. For a fixed number of excited neurons, the network frequency is ≈ 20 Hz lower in the GJ network than in the GJ-free network.

In Fig. 4E, the effect of the partial activation of the network on the synchrony index is shown. We find that for more than ≈ 10 excited neurons the GJ network is more synchronous than the GJ-free network.

In Fig. 4F, the influence of the number of excited neurons on the firing rate is depicted. Increasing the number of excited neurons decreases the firing rate from ≈ 280 Hz to ≈ 100 Hz for both networks. This decrease of the firing rate with growing number of active neurons can be explained by the fact that the feedforward, synaptic excitatory input per neuron is kept constant while the number of neurons receiving input is increased. So, increasing the number of active neurons is effectively shifting the inhibition-excitation balance to more recurrent, synaptic inhibition that, in turn, leads to lower firing rates.

In conclusion, we find that for an increasing number of active neurons the oscillation strength increases whereas the network frequency and the firing rate decrease. Additionally, GJs promote the oscillatory activity (larger oscillation strength), hence oscillations are possible at smaller numbers of active neurons (Fig. 4A&B). Moreover, GJs increase the synchrony in oscillating networks (Fig. 4E).

### 4.4 Delays of gap junction coupling potentials

Figures 2–4 have demonstrated that gap junctions between interneurons increase neuronal oscillation strength and synchrony. Synchrony, in turn, is strongly dependent on neuronal timing. Up to this point, we have assumed that gap junctions transfer their coupling potentials instantaneously. However, dendritic trees may cause delays of the GJ coupling potentials. Dendritic filtering should also affect amplitudes of GJ coupling potentials. To be able to assess the influence of delay and amplitude of GJ coupling potentials on network oscillations, an evaluation of the typical range of values is necessary.

To quantify GJ delays and amplitudes, we numerically simulate two GJ coupled multicompartmental neurons for variable GJ locations in the dendritic tree. Note that the coupling location is, for simplicity, always the same in both neurons (Fig. 5A). The neurons are coupled by a GJ that is modeled by an ohmic conductance of 1 nS. An action potential is generated in the presynaptic neuron, and the GJ coupling potential is measured in the soma of the postsynaptic neuron. In the following, we compare results from two different standard models of hippocampal PV+BCs (Saudargiene et al., 2015; Lee et al., 2014; see Methods for details).

**Figure 5:**
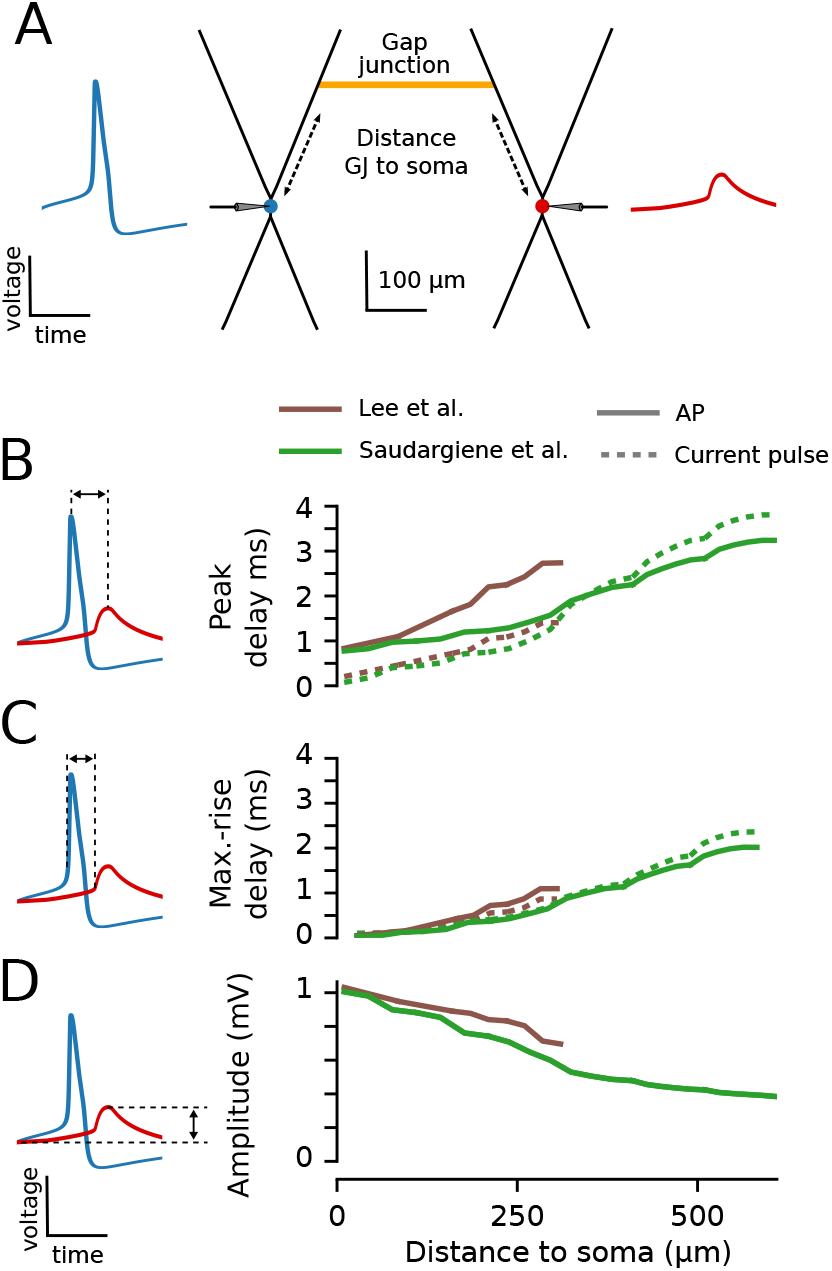
Delays and amplitudes of GJ coupling potentials in PV+BC multicompartment models. **A**, Schematic of two GJ coupled PV+BCs with simplified morphologies (model from Lee et al., 2014). AP (blue trace) in the left neuron (blue), and GJ coupling potential (red trace) in the right neuron (red). The position of the GJ is the same for both neurons. Thus, the distance that the AP has to travel from the soma to the GJ is the same as the distance the GJ coupling potential has to travel from the GJ to the soma. **B**, Left, The peak delay is calculated as the time between the maxima of the presynaptic AP (blue) and the postsynaptic GJ potential (red). Potentials not to scale. Right, peak delays for APs (solid lines) and waveforms evoked by short current pulses (dashed lines) for different values of GJ-soma distance as depicted in A (see Methods for details) for two different models (Lee et al. 2014; Saudargiene et al. 2015). The displayed delays in B correspond to GJ-coupling in the longer branches in the dendritic trees of the neurons in A, and delays are qualitatively similar for the shorter branches. B–D share the same x-axis. **C**, Same as B but for the maximal-rise (max.-rise) delay, i.e., the delay between the times of maximal rise of the potentials. **D**, Same as B and C but for the amplitude of the gap junction coupling potential. Here, only amplitudes of the action potential stimulus are displayed. For an overview of the parameters see Tables 1 and 2.

Results of simulations of two GJ coupled PV+BCs are depicted in Fig. 5B–D. We first calculated the peak delay, i.e., the delay from the peak of the presynaptic somatic AP of the first cell to the peak of the postsynaptic GJ potential in the soma of the second cell (Fig. 5B, left). This peak delay is calculated for different locations of the GJs in the dendritic tree, and hence different distances to the soma (Fig. 5B, right). The peak delay monotonically increases with increasing distance of the GJ from the soma. At the most distal ends of the dendrites, the measured delays are 2.7 ms and 3.2 ms for the Lee et al. (2014) and the Saudargiene et al. (2015) models, respectively (solid lines, Fig. 5B).

Surprisingly, even the peak delay at GJ locations close to the soma (< 50 μm) is > 0.5 ms. The main reason for this large minimum is that the peak delay is sensitive to the width of the presynaptic AP. Since the changes of the postsynaptic membrane potential (≲ 1 mV, see also Fig. 5D) are small in comparison to the amplitude of the action potential (≈ 100 mV), the width of the AP is a lower bound for the delay of the two peaks. Thus, the large delays for even short GJ–soma distances are the result of the broad action potentials of the two models (≈ 0.6 ms FWHM; see Methods), which are around double of what was measured in experiments (≈ 0.3 ms FWHM; Buhl et al., 1996; Kohus et al., 2016).

In essence, the delays at all distances, but most prominently at short distances, are overestimated in our simulations. This hypothesis is tested by replacing the presynaptic action potential by a fast bipolar current injection (FWHM< 0.1 ms) in a neuron model without active conductances in soma or dendrites. This change of the presynaptic pulse leads to very short peak delays at proximal GJ distances for both models ( ≈ 0.2 ms, dashed lines, Fig. 5B). For larger GJ-soma distances, we find that the signal generated by the current injection leads to longer delays than the AP for the Saudargiene et al. (2015) model. This behavior is caused by the lack of active conductances within the dendrites, which decelerates the transmission for longer distances.

Another way to remove the dependence of the delay on the width of the action potential is to measure the delay between the maximal rise of the AP and the maximal rise of the postsynaptic GJ potential. This maximal-rise delay is plotted in Fig. 5C. Resulting values are small (< 0.5 ms) at short distances (< 50 μm). The maximal-rise delay is also insensitive to the different presynaptic activations: action potentials and bipolar current pulses lead to similar delays. Furthermore, the estimated propagation speeds of signals within the dendritic tree, i.e., the slopes of the lines in Fig. 5B and C, are similar.

Finally, we measure the dependence of the amplitude of the GJ coupling potential on the location of the GJ (Fig. 5D). The amplitude of the postsynaptic GJ potential varies between 1.1–0.4 mV, and amplitudes monotonically decrease for increasing GJ–soma distance in both models. Amplitudes have to be treated with care because they also depend on the width of the presynaptic action potential, which is overestimated in the models used here. When we assume that the action potential would be half as wide, what is biologically plausible (Buhl et al., 1996; Kohus et al., 2016), and that the transferred current scales linearly with the width of the action potential, the amplitudes would be half as large. Explicitly, this scaling would lead to corrected GJ coupling potential amplitudes between ≈ 0.6 mV and ≈ 0.2 mV.

We conclude from the simulations that GJ delays in PV+BCs are short (≲ 0.5 ms) for proximal locations (≲ 100 μm) and can be quite long (> 1 ms) for distal locations (> 200 μm). While delays ≲ 0.5 ms were found in experiments (Galarreta and Hestrin, 1999; Tamás et al., 2000; Galarreta and Hestrin, 2001a), values > 1 ms have not been reported to the best of our knowledge. Further, simulations show that a presynaptic action potential elicits a gap junction coupling potential with an amplitude ≲ 0.6 mV, which is in agreement with experiments (≈ 0.5 mV, Tamás et al., 2000; ≈ 1 mV, estimated from Gibson et al., 1999; ≈ 0.5 mV estimated from Galarreta and Hestrin, 2001a).

### 4.5 Effect of gap junctions on ripple oscillations depends on gap junction delays

Having approximated the range of the delays for the GJ coupling, we can now analyze their effect on the steady-state ripple oscillations (Fig. 6). We vary the GJ delay *δ*_GJ_ from 0 to 2.4 ms, which shifts the point in time, when the active-spike component *β* increases the postsynaptic membrane potential (Eq. (1), Methods). In Fig. 3, we showed that the passive conductance *γ* has little effect on the network properties for zero delay, and hence we did not include nonzero delays in *γ* here. Otherwise we use the standard GJ parameters as in Figs. 1–4: *β* = 0.25 mV and *γ* = 1 nS. Furthermore, the GJ connection probability *p*_GJ_ is varied from 0 to 0.12 to provide a reference for the strength of the effect on the network properties that is caused by the delay.

**Figure 6:**
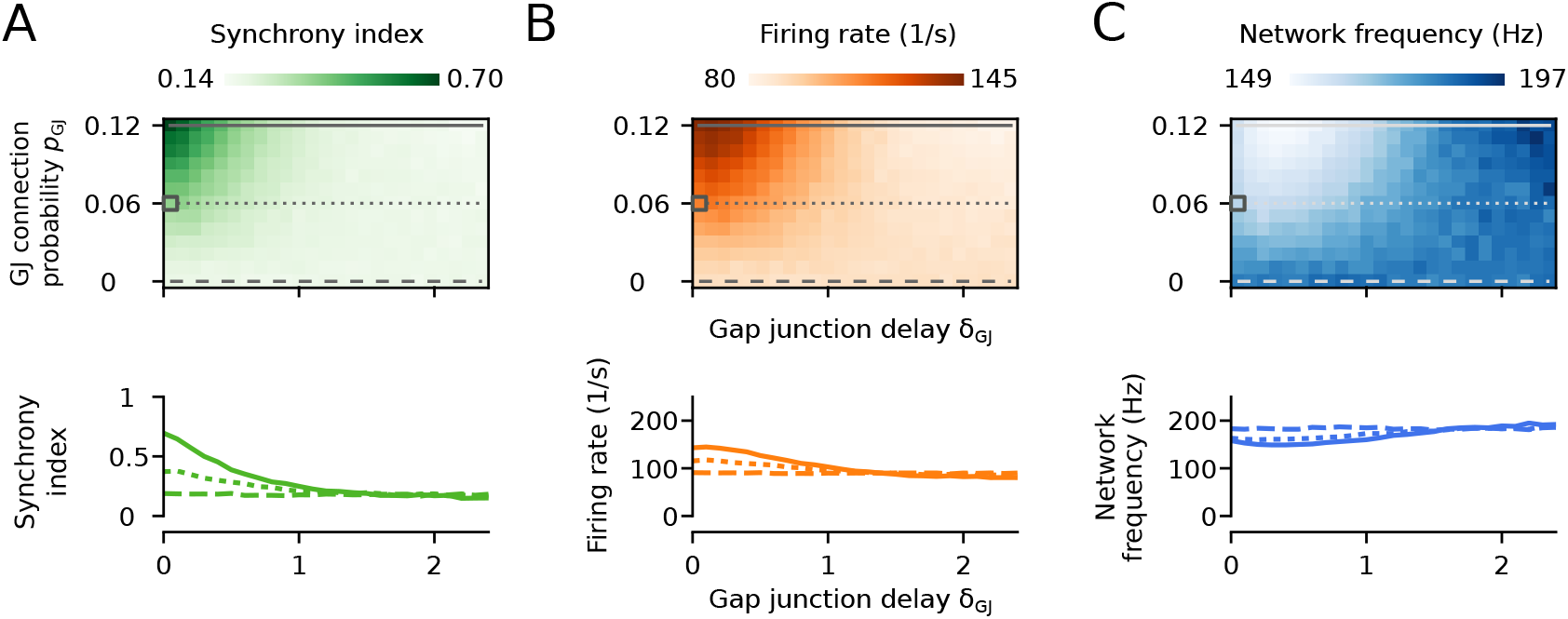
Small gap junction (GJ) delays only mildly decrease the impact of GJs on the network activity. Synchrony index (**A**), firing rate (**B**), and network frequency (**C**) as functions of the GJ connection probability *p*_GJ_ and the GJ delay *δ*_GJ_. The gray squares denote the standard parameters. Bottom row, Example traces at *p*_GJ_ fixed to 0, 0.06 and 0.12, as denoted by gray lines in the graphs at the top. Standard GJ parameters are used, i.e., *β* = 0.25 mV and *γ* = 1.0 nS. For an overview of the parameters see Tables 1 and 2.

The synchrony index is shown in Fig. 6A. We find an elevated level of synchrony in the network only for low values of the delay (≲ 1 ms). For longer delays (> 1.0 ms), the synchrony index is similar to the value in the GJ-free network (*p*_GJ_ = 0).

The neuronal firing rate varies from 80–145 Hz over the full range of the parameters (Fig. 6B). The firing rate is maximal at *δ*_GJ_ ≈ 0.1 ms, decreases for larger delays, and reaches the value of the GJ-free network at ≈ 1.3 ms.

The network frequency varies between ≈ 150 Hz and ≈ 200 Hz within the whole parameter range of *δ*_GJ_ and *p*_GJ_ (Fig. 6C). For high values of *p*_GJ_, the network frequency reaches its minimum at *δ*_GJ_ ≈ 0.3 ms. For *δ*_GJ_ > 1.3 ms the network frequency is at its reference value at *p*_GJ_ = 0.

Interestingly, we find the extremal values of the different network properties at different values of *δ*_GJ_: the firing rate and the network frequency reach their extremal values at ≈ 0.1 ms and ≈ 0.3 ms, respectively.

In conclusion, GJ potentials have to fall into a narrow time window (≲ 0.5 ms) of the oscillation cycle, at which the average membrane potential is close to spiking threshold, to promote synchrony, increase the firing rates, and decrease the network frequency. This temporal window is short since it is only a fraction of the total period (≈ 5 ms) of the ripple oscillations.

## 5 Discussion

Parvalbumin-positive basket cells (PV+BCs) are coupled by gap junctions (GJs; Katsumaru et al. 1988; Fukuda and Kosaka 2000a; Galarreta and Hestrin 2001a), and in the hippocampal area CA1 they form recurrently coupled interneuron networks (INT-INT) that are hypothesized to generate ripples (Ylinen et al., 1995; Klausberger et al., 2003). To test the functional relevance of these GJs for ripple oscillations, we used a biologically plausible network model of PV+BCs that reproduced ripple oscillations under transient and steady-state input (Donoso et al., 2018).

Our simulations showed that interneuronal GJs, especially action potentials transmitted by GJs, increase the synchrony and the mean firing rate of the interneuronal network during ripple oscillations, but only mildly decrease the frequency of the ripple oscillations. Furthermore, GJs reduce the minimum number of active interneurons required for ripple oscillations in such INT-INT networks. Finally, GJ transmission delays can vary from 0 to ≈ 3 ms, which depends on the somatodendritic GJ location and on the PV+BC model. We demonstrated that only small GJ delays (≲ 0.5 ms) promote ripple oscillations.

We predict that hippocampal ripple oscillations that are generated by INT-INT networks are affected by deactivation of the interneuronal GJs (Ylinen et al., 1995). To test this hypothesis, we propose to record the spiking activity of CA1 hippocampal PV+BCs, e.g., by extracellular multi-electrode recordings or by intracellular recordings of membrane potentials, while the properties of gap junctions among PV+BCs are selectively altered.

### Experimental evidence for the function of gap junctions in ripple oscillations

Many studies have already tried to test the functional relevance of GJs for hippocampal ripple oscillations, using either chemical GJ blockers (Ylinen et al., 1995; Draguhn et al., 1998; Hormuzdi et al., 2001; Maier et al., 2003; Pais et al., 2003; Traub et al., 2003; Buhl et al., 2003; D’Antuono et al., 2005; Behrens et al., 2011) or connexin36 knockout (Cx36KO) mice (Hormuzdi et al., 2001; Maier et al., 2002; Pais et al., 2003; Buhl et al., 2003) lacking the GJ protein Cx36, which has been found in pyramidal cells (Condorelli et al., 2000) and interneurons (Venance et al., 2000). Most studies that rely on the chemical GJ blockers octanol, carbenoxolone, or halothane (Ylinen et al., 1995; Draguhn et al., 1998; Hormuzdi et al., 2001; Maier et al., 2003; Pais et al., 2003; Traub et al., 2003; Buhl et al., 2003; Behrens et al., 2011) find a strong suppression or abolishment of SWRs (but cf. D’Antuono et al., 2005), and, hence, do not allow conclusions about changes of the frequency of ripple oscillations. These findings are contrasted by experiments using Cx36KO mice (Hormuzdi et al., 2001; Pais et al., 2003; Buhl et al., 2003) or the GJ blocker mefloquine (Behrens et al., 2011) that only find mild effects on SWRs. In the studies in which ripple oscillations were still observed after a putative GJ block, the ripple frequency was not affected (Maier et al., 2003; Buhl et al., 2003; D’Antuono et al., 2005; Behrens et al., 2011 but cf. Maier et al., 2002).

These contradictory results might be explained by several confounding factors: GJ blockers are not specific and have strong side-effects (Juszczak and Swiergiel, 2009), SWRs were stimulated by different means (GABA, Traub et al., 2003; kainate, Hormuzdi et al., 2001; Pais et al., 2003; picrotoxin, D’Antuono et al., 2005, KCl, Nimmrich et al., 2005, Ca^2+^ - free ACSF, Hormuzdi et al., 2001), and networks of Cx36KO might be altered due to compensatory effects during development. Moreover, GJ blocker and Cx36KO experiments are not specific for GJs between PV+BCs but also interfere with putative GJs between pyramidal neurons.

GJs between pyramidal cells are the major element of an alternative theory for the origin of ripple oscillations (Traub et al., 1999), albeit evidence for pyramidal GJs is sparse for mature pyramidal cells (Rash et al., 1997; Condorelli et al., 2000; Mercer et al., 2006; Wang et al., 2010). According to the hypothesis from Traub et al. (1999) a block of GJs between pyramidal cells would abolish ripple oscillations, which is in contrast to experiments that observed only mild effects on the network dynamics (Hormuzdi et al., 2001; Pais et al., 2003; Buhl et al., 2003; Behrens et al., 2011).

### How many interneurons are necessary to generate ripple oscillations?

Our simulations showed that GJs decrease the minimal number of excited interneurons that is required to generate ripple oscillations (Fig. 4), and the minimum number is on the order of tens of neurons. For a sufficiently large number of interneurons, ripple oscillations can be more robustly generated when GJs are present. Our estimates depend on a specific set of parameters, yet, the qualitative observations that GJs decrease the number of necessary neurons was true for all tested sets of biologically plausible parameter ranges.

Some experimental constraints for the minimum number of interneurons required for ripple-like oscillations were obtained via optogenetics. Schlingloff et al. (2014) found that the activation of ≈ 150 PV+BCs in hippocampal CA3 slices was enough to generate ripple-like steady-state oscillations. While this supports the INT-INT hypothesis, it was challenged by the in vivo study by Stark et al. (2014) in CA1, where the optogenetic excitation of ≲ 20 PV+BCs was not sufficient to generate ripple-like activity. In the light of our findings, we argue that the number of directly activated interneurons by Stark et al. (2014) was still below the threshold of neurons required for ripple-like oscillations, and consequently the results by Stark et al. (2014) do not necessarily reject the INT-INT hypothesis for ripples.

To test our prediction that a certain minimal number of PV+BCs is necessary for ripple oscillations, we propose the following experiment. The EFP is recorded in a hippocampal slice during an optogenetic direct activation of a variable number of PV+BCs. There are three specific predictions: First, ripple-like oscillations require the activation of more than a sufficient (minimal) number of PV+BCs. Second, this minimum is smaller for wild-type mice with intact GJs in comparison to Cx36KO mice. Third, the firing rate and the network frequency decrease with an increasing number of activated neurons.

### Gap junction transmission delays

We calculated the delays of GJ potentials between hippocampal PV+BCs to be in the range 0–3 ms, which depends on the GJ location in the dendritic tree (Fig. 5) and on the specific neuron model (Lee et al., 2014; Saudargiene et al., 2015). In contrast, only delays below ≲ 0.5 ms have been found experimentally (Galarreta and Hestrin, 1999; Tamás et al., 2000; Galarreta and Hestrin, 2001a). Such short delays require proximal GJ coupling (≲ 100 μm from soma; Fig. 5) as observed between neocortical PV+BCs in ultrastructural studies (Tamás et al., 2000). Our analysis showed that with such small delays GJs can still promote ripple oscillations (Fig. 6). Conversely, if the GJs are located more distally, as found in ultrastructural studies in the hippocampus (> 200 μm; Fukuda and Kosaka, 2000b), the delays of the GJ potentials were long (> 1 ms). In our network simulations, we found that GJs with such long delays do not promote ripple oscillations, possibly because the GJ potentials are outside of the time window within each oscillation cycle in which neurons are spiking.

Motivated by our simulations (Fig. 5) and experimental evidence of distal GJ coupling (Fukuda and Kosaka, 2000b), we predict that GJ potentials with long delays (> 1 ms) do exist between PV+BCs (Fig. 6). Such long delays have not been observed in experiments to the best our knowledge. Moreover, we predict fast GJ potentials (≲ 0.5 ms) between hippocampal PV+BCs in CA1 (Bartos et al., 2002) in analogy to neocortical findings (Galarreta and Hestrin, 2001a) because proximal GJ coupling (≲ 100 μm) of PV+ interneurons was shown in ultrastructural studies (Fukuda and Kosaka, 2000b).

### Limitations of this study

Our network simulations are based on single-compartment leaky integrate-and-fire neurons, which is a major simplification of the neuronal dynamics because such models do not include action potentials. To be able to include GJs in this network model and to simulated coupling potentials evoked by action potentials, we used the two-parameter model for the GJs by Lewis and Rinzel (2003). Further, the neurons in our network model do not have any physical extension, and hence cannot describe propagation of signals within the neurons. To account for such delays, we included them in electrical and chemical couplings.

### Comparison to other theoretical studies

We found that GJs increase the synchrony of neuronal oscillations. This confirms results of previous approaches that either employed analytical methods on idealized networks (Lewis and Rinzel, 2003; Kopell and Ermentrout, 2004) or computational methods on more biologically plausible networks (Traub et al., 2001; Bartos et al., 2002; Maex and De Schutter, 2003; Guo et al., 2012). While all of these studies showed that GJs can increase synchrony for various network settings, none of these studies had a focus on hippocampal ripple oscillations nor considered different strengths and connectivities of GJ coupling. We showed that GJs increase the synchrony of ripple oscillations for a large range of realistic GJ parameters. Further, we provided explicit predictions for the role of interneuronal GJs in ripple oscillations, and systematically studied the size and impact of GJ delays, which has only been considered implicitly (Traub et al., 2001; Maex and De Schutter, 2003) or has been neglected (Bartos et al., 2002; Guo et al., 2012) in previous studies.

### Conclusion

GJs between PV+BCs promote and stabilize hippocampal ripple oscillations if the GJ delay is ≲ 0.5 ms. We find that for such short delays the GJ coupling has to be proximal (≲ 100 μm from soma). We confirm that such interneuronal GJs have a weak effect on the ripple frequency, and that they are not the primary pacemaker of the ripple oscillations. These findings support the INT-INT hypothesis of ripple oscillations, which assumes that recurrent chemical connections of interneurons set the oscillation frequency.

